# UVB induces meibomian gland dysfunction and ocular surface damage in rat mimicking ocular rosacea

**DOI:** 10.1101/2025.03.31.646002

**Authors:** Linxin Zhu, Daniela Rodrigues-Braz, Emmanuel Gélizé, Marine Crépin, Caroline Peltier, Coralie Lheure, Xavier Morel, Sandrine Moret, Jean-Louis Bourges, Olivier Berdeaux, Francine Behar-Cohen, Min Zhao

## Abstract

Ocular Rosacea (OR) is a common chronic inflammatory disease that affects the ocular surface and eyelids. Meibomian gland dysfunction (MGD), often associated with OR, can lead to dry eye syndrome and vision-threatening corneal complications. OR is underdiagnosed and incurable, highlighting the need for a deeper understanding of the pathogenesis of MGD associated with OR to develop targeted treatments. In this study, rat upper eyelids were exposed to UVB, a known triggering factor for rosacea, for five days. UVB exposure caused acute damage to the eyelid skin and meibomian glands (MG) together with increased oxidative stress, mitochondrial dysfunction, apoptosis, inflammation, and elevated lipid production by day 5. During the two-week healing process, inflammation and lipid hyperproduction tended to normalize, but hyperkeratinization of MG ducts persisted, and meibocyte stem cells continued to decrease. This led to MGD and was likely responsible for corneal epithelial defects observed on day 19. The progressive fibrosis in rat MG was similar to that observed in MG biopsy tissues from OR patients, suggesting that UVB causes chronic and irreversible damage to MGs. Transcriptomic intersection analysis revealed shared gene regulation patterns between UVB-induced changes in rat MG and human rosacea and MGD. Additionally, UVB altered Meibum lipid composition in ways that mimic human MGD. This model provides a valuable tool for studying the pathophysiology of MGD associated with OR and for evaluating potential treatments.

## Introduction

Rosacea is a common chronic cutaneous inflammatory and neurovascular disease that affects around 5.46% of the adult population [1]. Ocular manifestations occur in up to three quarters of the total rosacea population [2–5]. It is characterized by eyelid margin telangiectasia, blepharitis, keratitis and conjunctivitis with complaints of photophobia, blurred vision. Meibomian gland dysfunction (MGD) is associated with up to 82% of patients with ocular rosacea (OR) [6]. Structural and functional alterations of MG modify the quantity and lipid composition of meibum, causing the dry eye symptoms associated with OR [7]. Around one third patients suffering from OR develop corneal pathology ranging from mild inferior punctate epithelial defects to potentially sight-threatening corneal neovascularization, scarring, corneal thinning, ulceration and perforation in the most severe cases [8]. OR may precede, follow, or occur simultaneously with the skin changes. Diagnosis of OR is largely based on clinical signs, but clinical features are generally not specific to OR, as similar ocular manifestations can be presented in other ophthalmic diseases [9], making OR often underdiagnosed. OR is incurable. Current therapeutic options only offer indirect, nonspecific approaches or address already existing ocular surface damages.

The pathophysiology of OR is not fully understood, involving genetic predisposition, dysregulation of the innate and adaptive immune system, vascular and neuronal dysfunction, and microorganisms such as *Demodex folliculorum* [10–12]. Activation of the toll-like receptor (TLR) 2 [13] and transient receptor potential (TRP) ion channels [14] with subsequent release of inflammatory mediators from multiple cell types including keratinocytes, mast cells, neurons, endothelial cells, macrophages, fibroblasts and Th1/Th17 cells has been considered as primary or secondary causative mechanisms of rosacea [10]. Triggering factors such as heat, psychosocial stress, ultraviolet (UV) light, spicy food, smoking and alcohol can initiate the onset of symptoms and exacerbate them but sun exposure is the most recognized aggravating factor [15].

Solar UV radiation is composed of UVA, -B and -C, which have different electro-physical properties. UVB (290-320 nm) can induce harmful effects on DNA and cells in the skin and eye surface [16,17] since the cornea, conjunctiva and eyelids absorb more than 90% of UVB below 300 nm. Acute overexposure to UVB typically results in photokeratitis characterized by severe pain, photophobia, tearing and inflammation. Chronic UVB radiation is an important risk factor for cataract and pterygium but can also cause climatic droplet keratopathy and ocular surface squamous neoplasia [16,18,19]. UVB radiation may contribute to the development of skin and ocular rosacea by causing oxidative damage and inducing pro-inflammatory cytokines (IL-1α, IL-1β, IL-6, IL-8, IL-17), chemokine (C-C motif) ligand 5 (CCL5), tumor necrosis factor-α (TNF-α), as well as vasoactive (VEGF) and neuroactive factors [10,20–22].

Appropriate animal models are key for unveiling the mechanisms of human diseases and for therapeutic development. Murine models for skin rosacea include the cathelicidin LL37 intradermal injection model which has become the most intensively used model for skin rosacea research, KLK-5-induced inflammation, croton oil-induced polymorphonuclear leukocyte-mediated skin inflammation, 12-O-Tetradecanoylphorbol-13-acetate inflammation model, arachidonic acid inflammation model, and UVB-induced erythema models [23]. In UVB models, animals were exposed either to a high-dose of UVB light (120 mJ/cm^2^) [24] or to UVB for 5 minutes every other day for 10 days and then daily from day 11 to day 14 [21]. These two experimental conditions induce vascular reactions and angiogenesis in mouse skin mimicking erythematous rosacea. None of these models develop OR, and the MGD associated with OR has never been studied.

Various genetic mouse models have been developed and used to evaluate the contribution of different molecular pathways and cell types to MGD [23]. Some mice also develop age-related MGD similar to human disease, with MG atrophy already evident at 1 year of age [25]. But none of these models recapitulate the complete pathophysiological process of MGD associated with OR.

In this study, five days of UVB irradiation of rat eyelids resulted in ocular surface signs similar to those seen in human OR. UVB induced hyperkeratinization and inflammatory cell infiltration in the epidermis and MG, fibrosis in the interstitial tissue, meibocyte dysfunction, apoptosis, mitochondrial dysfunction and oxidative stress in the MG. Two weeks post-UVB exposure, corneal epithelial defects and barrier disruption were observed as a consequence of MGD. Transcriptomic analysis identified UVB-induced functional gene sets and differentially expressed genes shared with human MGD and rosacea. Lipidomic analysis also revealed changes in lipid composition within the MG.

## Materials and Methods

### Human eyelid samples collection

The collection and storage of human biological samples were approved by local ethics committee CCP Ile de France 1 (no. 2016-nov-14390). Eyelid biopsy samples were obtained from 2 patients with OR (1 man and 1 woman) and 3 control subjects without rosacea (from tarsal surgery) (2 man and 1 woman), age ranged between 40 and 90 years. Donors provided written consent for further use of samples in research.

Tissues were fixed in 10% formalin for 24 hours. After dehydration, tissue was embedded in paraffin wax and cut into 7 um thick slices for histological staining.

### Animal Experiments

All experiments were performed in accordance with the European Communities Council Directive 86/609/EEC and French national regulations and approved by local ethics committee (# 23478-2020010317557546 v4, Charles Darwin). Adult female Sprague-Dawley rats (12 weeks, 450-500 g, Janvier Labs, Le Genest-Saint-Isle, France) were used for UVB irradiation of eyelids. All animals were kept in pathogen-free conditions with free access to food and water and housed in temperature-controlled room with a 12-h light/12-h dark cycle. Anesthesia was induced by intraperitoneal ketamine 100 mg/kg and Xylazine 10 mg/kg. Animal were euthanized by intraperitoneal injection of fatal dose of Euthasol® Vet.

### UVB irradiation on rat eyelids

The upper eyelids of rats were shaved and exposed to 1000 µW/cm^2^ of UVB (312 nm, Herolab, Wiesioch, Germany) for 5 min (300 mJ/cm^2^) daily for 5 days, protocol adapted from previous studies [26] concerning the different ocular tissues. The cornea and the whole body were covered and protected (Figure 1A). Lesions of the eyelid skin and eyelid margin were photographed under the microscopy at D0, D3 and D5 (Figure 1B). Rats were euthanized after 5 days of irradiation to evaluate the acute damage of ocular surface tissues (n = 8 rats). Rats without UVB irradiation serve as controls (n = 7 rats).

**Figure 1.**
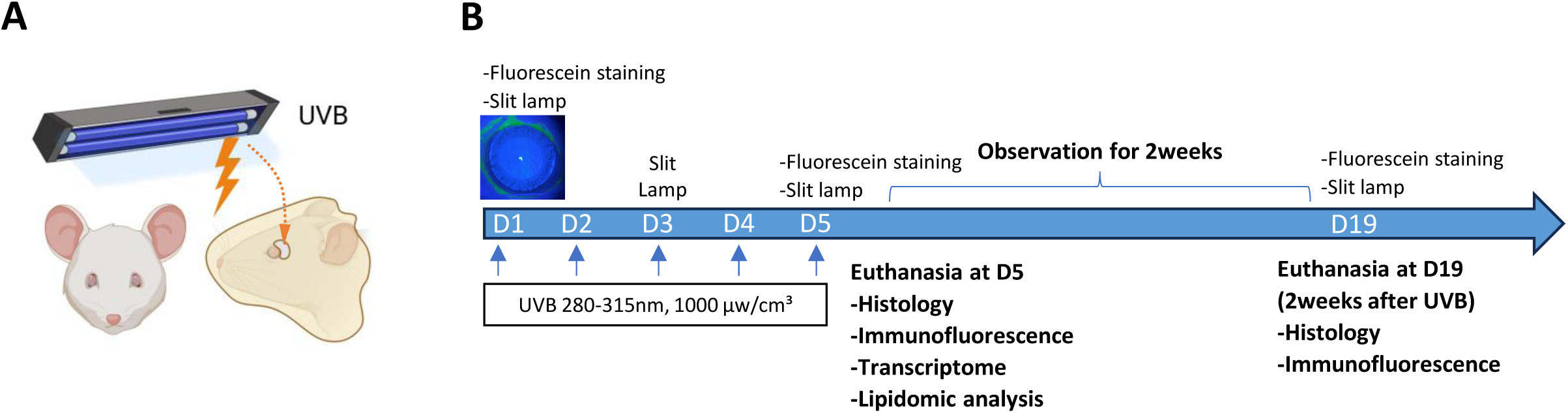
Schematic representation of the UVB irradiation procedure on rat eyelids. **A.** Rat upper eyelids are exposed to UVB. The cornea and the rest of the body are covered. **B.** Experimental design.

To assess the prolonged effects of UVB irradiation on meibomian gland function and on the cornea, a group of rats (n = 7) were followed up until 2 weeks after UVB exposure (Figure 1B). Corneal fluorescein staining was performed before and after 5 days UVB irradiation and 2 weeks after UVB exposure. Corneal epithelial defect was recorded under the slit-lamp microscope with a cobalt blue filter. Rats were then euthanized, eyelids and corneas were dissected for morphological and immunohistochemical analysis (Figure 1B). Corneal defects were graded according to the criteria previously described [27,28]. In brief, the cornea was divided into 4 quadrants, individual scores were made for each quadrant respectively: 0, absent; 1, slightly punctate staining less than 30 spots; 2, punctate staining more than 30 spots but not diffuse; 3, severe diffuse staining without positive plaque; and 4, positive fluorescein plaque. The 4 scores for each quadrant were summed to reach a final score for each cornea sample, with a possible total score of 16 points. Comparison was conducted among different time points.

### Hematoxylin and eosin stain

Human and rat eyelid paraffin sections were dewaxed in xylene and rehydrated in a series of decreasing concentrations of ethanol. After staining with hematoxylin reagent (RAL Diagnostics, #320550-2500, Bordeaux, France) for 15 mins, slides were rinsed by tap water to remove floating color. Then the sections were immersed in eosin solution (RAL Diagnostics, #312730-0100, Bordeaux, France) for 4 minutes and rinsed again by tap water. Following sequential baths of absolute ethanol and xylene, tissue sections were mounted by Eukitt® (O.Kindler GmbH, #200080, Freiburg, Germany). Morphological changes were observed under microscope (Olympus BX51, Rungis, France) equipped with a CCD camera (Olympus DP70).

### Sirius red staining

For collagen observation, after dewaxing and rehydration, human and rat eyelid sections were incubated with 1% Sirius red in picric acid for 15 minutes. After different baths of acetic acid and absolute ethanol, slides were mounted and observed under microscope. The content of collagen fibers was quantified using Fiji ImageJ software (version 1.54b, Wayne Rasband, National Institutes of Health, Bethesda, MD, USA) by performing a binary conversion of the images on red channel to quantify the positive % of area occupied by these targets. The density of acinar units was quantified by Fiji ImageJ software using angiogenesis analysis, and the number of complete meshes were calculated and compared. All quantifications were normalized to the area of the tissue analyzed.

### Immunohistochemistry

Rat eyelid paraffin slides were used for immunohistochemical staining of markers for oxidative stress. After dewaxing and rehydration, eyelid sections were incubated with 10 mM citrate buffer pH 6.0 for 20 min at 100 °C for antigen retrieval. The endogenous peroxidase was blocked with 3% hydrogen peroxide for 30 min, followed by incubation with TNB blocking buffer reconstitute with blocking reagent powder (AKOYA Biosciences, #SKU FP1012, Massachusetts, USA) for 30 min. Primary antibodies, rabbit anti-Nitrotyrosine (NT, 1:200, Thermo Fisher Scientific, #BS-8551R, Saint Aubin, France), and rabbit anti-4 Hydroxynonenal antibody (4-HNE, 1:200, Abcam, #ab46545, Cambridge, UK) were incubated for overnight at 4 °C. After 3 washes with PBST, tissue sections were then incubated with biotinylated goat anti-rabbit secondary antibody (1:500, Vector Laboratories, #BA-1000, Eurobio Scientific, Les Ulis, France) for 1h at room temperature, followed by Vectastain Elite ABC reagent (Vector Laboratories) for 30 min. The reaction was accomplished with 3,3’- diaminobenzidine (DAB) substrate and stopped in 50 mM TRIS pH 7.6 for 10 min. Sections were then mounted with Eukitt®. Negative controls were performed without primary antibody.

### Immunofluorescence

For immunofluorescence staining, rat eyelid and corneal cryosections were fixed in 4% paraformaldehyde (PFA) for 15 min, followed by 3 washes of PBS for 5 min each. Sections were then permeabilized with 0.1% Triton X-100 in PBS for 30 min and blocked with 5% goat serum in PBS for 30 min. Tissues were then incubated with primary antibodies for overnight at 4 degrees. Following 3 washes of PBS, sections were further incubated with secondary antibodies for 1h at room temperature. Slides were then stained for 5 min with 4’,6-Diamidino-2-Phenyl-Indole (DAPI; 1: 5000) and mounted with gel mount (Dako, Agilent, Les Ulis, France). Negative controls were performed without the primary antibody. Sections were then observed by Olympus fluorescence microscope (Olympus BX51). Primary antibodies and dilutions used in this study include: rabbit anti-P63α (1:100, Cell Signaling Technology, #4892, Ozyme, Saint-Cry-L’Ecole, France), rabbit anti-PPARγ (1:100, Cell Signaling Technology, #2435S), mouse anti Ki67 (1:100, Cell Signaling Technology, #9449 (8D5), Massachusetts, USA), rabbit anti Cytokeratin 1 (1:500, Abcam, #ab185628), rabbit anti Cytokeratin 10 (1:500, Abcam, #ab76318), rabbit anti-IBA1 (1:400, Wako #019-19741, Richmond, VA, USA), mouse anti-ED1 (1:200, Bio-Rad #MCA341R, Colmar, France), rabbit anti TIM23 (1:100, Proteintech #11123-1-AP, Planegg-Martinsried, Germany), rabbit anti TOM20 (1:400, Proteintech #11802-1-AP), mouse anti E-Cadherin (1:400, Abcam #ab231303), rabbit anti ZO-1 (1:200, Thermo Fisher Scientific, #40-2200,). Secondary antibodies used were Alexa Fluor 488-conjugated goat anti-rabbit IgG (1:200; Thermo Fisher Scientific, #A11008), Alexa Fluor 488-conjugated donkey anti-mouse IgG (1:200; Thermo Fisher Scientific, #A21202), Alexa Fluor 594-conjugated goat anti-rabbit IgG (1:200, Thermo Fischer Scientific, #A11012) and Alexa Fluor 594-conjugated donkey anti-mouse IgG (1:200, Thermo Fischer Scientific, #21203). For further quantification, the ratio of P63-, PPARy- and Ki67-positive cells to all DAPI-stained cells in whole gland tissue was calculated respectively; the expression of TIM23 and TOM20 was quantified according to their average fluorescence intensity in whole gland tissue; all data were expressed as relative values compared to the control group. The number of IBA1- and ED1-positive cells were counted per field.

### Nile red staining

For lipid staining, fresh frozen rat meibomian gland sections were incubated with 1µg/ml Nile red (Sigma-Aldrich, #72458, Missouri, USA) in PBS for 1 hour at 4 degree. Tissues were then fixed in 4% PFA for 15 min and followed by DAPI staining and PBS washes. Images of staining were observed and taken by confocal microscope (Zeiss LSM 710, Oberkochen, Germany). Nile red-stained lipid droplets were quantified using Fiji imageJ software by calculating the percentage of the stained area relative to the total analyzed area.

### TUNEL assay

Cell apoptosis was evaluated by TUNEL Assay according to manufacturer’s instruction (Roche Diagnostics, Mannheim, Germany). Nuclei were counterstained.

### Statistics Analysis

Data are expressed as mean ± SD. Statistical analysis was performed using the GraphPad Prism (GraphPad Software, version 9, San Diego, CA, USA). The Mann– Whitney test was used to compare two groups. The Kruskal–Wallis test followed by Dunn’s test was used to compare more than two groups. The Friedman test was used to compare more than 2 matched groups. A p-value less than 0.05 was considered statistically significant.

### Transcriptomic analysis of rat meibomian gland

After 5 days exposure to UVB, UVB-irradiated rats (n=3) and their age- and sex-matched non-irradiated control rats (n=4) were euthanized. Upper eyelids were removed and meibomian gland tissues were isolated and snap frozen in liquid nitrogen. RNA extraction was performed using the Qiagen RNeasy Kit (Cat. 74004, Qiagen©, Hombrechitikon, Switzerland), and 1 µg of extracted RNA was sent for RNA sequencing at the iGenSeq transcriptomic platform of the Brain and Spine Institute (ICM, Paris, France). RNA quality was checked by capillary electrophoresis and RNA integrity numbers (RIN) ranging from 8.8 to 9.3 was accepted for library generation. Quality of raw data was assessed using FastQC. Poor quality sequences and adapters were trimmed or removed with fastp, using default parameters, to retain only good quality paired reads. Illumina DRAGEN bio-IT Plateform (v3.8.4) was used for mapping on rn7 reference genome. Library orientation, library composition and coverage along transcripts were checked with Picard tools. The transcriptomic data were analyzed on the online platform of the Paris Brain institute (http://quby.icm-institute.org). Firstly, differential expression analysis using DESeq2 method was utilized to identify the differentially expressed genes (adjusted p-values, pFDR). FDR threshold was set at 0.05, log2 fold-change threshold was set at 0.5. Then, an enrichment analysis was performed (by over-representation analysis) in different gene-set collections including Reactome gene sets (Reactome subset of Canonical pathways) and Gene Ontology gene sets (C5:GO) and Hallmark gene sets (H). Further intersection analyses were conducted between the transcriptomics of MGs from patients with MGD (GSE17822) and from rats in our study, as well as between the transcriptomics of facial biopsies from rosacea patients (GSE65914) and rat MGs in this study.

### Lipidomic analysis of rat meibomian gland

Meibomian gland tissues from UVB-irradiated (n = 11) and non-irradiated rats (n = 19) were used for lipidomic analysis. Lipids were extracted using the Folch method [29]. The lipid-containing phase was collected and dried under a nitrogen stream. The pellet was reconstituted in 200 µL of CHCl3/MeOH (2:1).

Lipid separation was performed on a reversed-phase C18 column (RPC) and a hydrophilic interaction liquid chromatography (HILIC) column as previously described [29]. The injection volume was 5 µL. For RPC, the mobile phase A consisted of acetonitrile/water (60:40, v/v) and phase B consisted of isopropanol/acetonitrile (90:10, v/v), both containing 10 mM ammonium formate and 0.1% formic acid. The solvent-gradient system was as follows: 0 min 60% (A), 10 min 100% (A), 12 min 100% (A), 12.1 min 40% (A), and 17 min 40% (A). For HILIC, the mobile phase A consisted of acetonitrile/water (95:5, v/v) and phase B consisted of acetonitrile/water (50:50, v/v), with 10 mM ammonium acetate in both phases. The solvent-gradient system was as follows: 0 min 100% (A), 5 min 80% (A), 6.5 min 60% (A), 8 min 60% (A), 8.1 min 100% (A), and 10 min 100% (A).

Lipidomic analysis was performed using a Thermo Ultimate^TM^ 3000 coupled to an Orbitrap Fusion^TM^ Tribid Mass Spectrometer (MS) equipped with an EASY-MAX NG^TM^ Ion Source (Thermo Scientific, San José, CA, USA). MS acquisition in full scan was performed in positive and negative modes, with a static spray voltage at 3,500 V and 2,800 V, respectively. The detector used was an Orbitrap with a resolution of 120,000, on a m/z range of 200-1200 using quadrupole isolation. MS were recorded using a maximum injection time of 50 ms, a standard automatic gain control target, a radio frequency lens at 50% - excepted with the RPC column in positive mode (60%)- and in centroid mode. Source parameters were set as follows for RPC column: ion transfer tube temperature of 285°C, vaporiser temperature of 50°C, sheath gas flow rate of 63 arbitrary unit (au), sweep gas of 20 au, and auxilary gas flow rate of 25 au. Source parameters for HILIC were set as previously described [29].

For lipidomic data acquisition, the Orbitrap Fusion was controlled by XcaliburTM 4.3.73.11 software (Thermo Scientific). Intensities values were extracted with LCMS R package [30], using a homemade database containing 2105 lipid species from 13 classes – created according to data found in literature and in biological samples, and then checked with FreestyleTM 1.5 software (Thermo Scientific). The analysis of the UVB or non-UVB samples were conducted during two batches.

Regarding the data analysis, the same procedure was applied to HILIC and C18 data. First, the intensities were pre-processed as follows: (i) intensities lower than 1000 were considered as noise and replaced by missing values, (ii) lipid species with more than 90% missing values were removed from the analysis, (iii) the signal was normalized per sample, and (iv) the batch effect was removed. Then, for both positive and negative modes, the same approach was used. To begin with, an exploratory analysis (PCA) was conducted to visualize potential groups or outliers and to determine if the UVB factor appeared as the primary source of variability in the data. Next, Mann-Whitney Wilcoxon tests were conducted to detect the UVB effect in all lipid species. The p-values were corrected using the Benjamini-Hochberg method (BH or FDR), and volcano plots were generated. Finally, an approach considering lipid classes instead of lipid species was applied using the same methods (PCA, followed by Mann-Whitney Wilcoxon tests).

## Results

### Eyelids from OR patients exhibited inflammatory cell infiltration and fibrosis in the MGs

To identify morphological changes in the MGs of OR patients, eyelid biopsy tissues from 2 OR patients and 3 control subjects were used for histological staining. As shown in Figure 2A, abundant inflammatory cellular infiltration was observed among acini in the MG, beneath the epidermis and around the hair follicles of eyelid tissues from OR patients compared to control subjects. In OR patients, MG acini appeared more compact and irregular, whereas MG acini in control subjects were well-organized. Sirius Red selectively staining interstitial collagen fibers showed enhanced staining in MGs of OR patients compared to controls, indicating fibrosis (Figure 2B).

**Figure 2.**
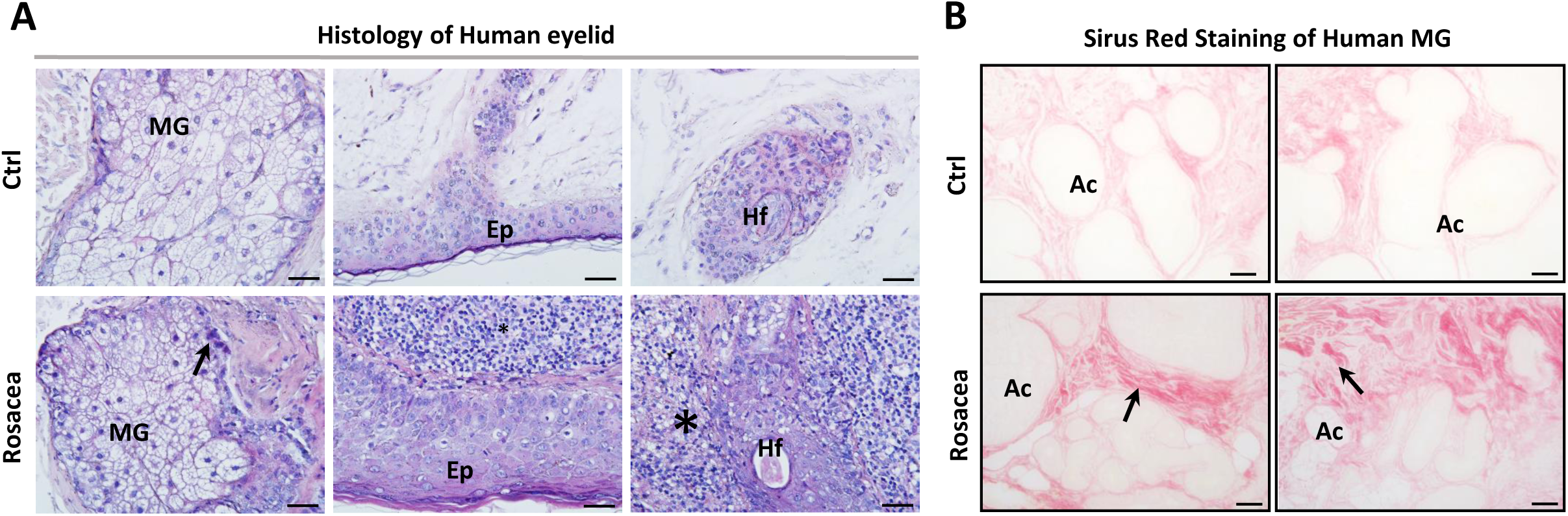
Histological changes in the meibomian glands of patients with ocular rosacea. **A.** HE staining of human meibomian glands (MG) shows inflammatory cell infiltration around acini (Ac, arrow), beneath the epidermis (Ep) and around hair follicles (Hf) (asterisks) of eyelid biopsy tissues from patients with ocular rosacea. The acini appear smaller and more compact than those of control subjects (Ctrl). Scale bar: 50um. **B.** Sirius Red shows intense collagen staining in MGs of rosacea patients compared to control subjects (arrows). Scale bar: 50um.

### UVB-irradiated rat eyelids presented similar clinical signs and histological features as in OR patients

Slit-lamp revealed redness and swelling of rat eyelid skin, telangiectasia and hyperemia at the lid margin and eyelash drop out on days 3 to 5 of UVB irradiation (Figure 3A). After 5 days of UVB irradiation, the eyelid skin was covered with a yellowish exudate (Figure 3A). In addition, swelling of the conjunctiva and MGs was observed on the inner surface of the eyelid (Figure 3A, black and white picture). Histology of rat MGs from irradiated eyelids showed cellular infiltration among the acini, around the gland ducts and beneath the epidermis (Figure 3B). Morphological abnormalities in MGs included irregular epithelial layout, cell shedding and inflammatory cell exudation in gland ducts, cavitation and nuclei distortion in acinar cells (Figure 3B). In UVB-irradiated eyelids, Sirius Red staining increased significantly on day 5 (29.81 ± 1.69% vs. 25.28 ± 0.96% in the control group, n = 5 rats in both groups, p = 0.0005) and further increased on day 19, two weeks after UVB exposure (32.15 ± 1.34%, n = 5 rats, p = 0.0458 vs. UVB group, p < 0.0001 vs. control group), suggesting fibrosis (Figure 3C and 3D). The number of acini per field increased to 128.3 ± 11.62 after 5 days of UVB, vs. 80.7 ± 11.11 in the control group (n = 5 rats, p = 0.0006). Two weeks after UVB exposure, the number of acini was 143.5 ± 19.22 (n = 5 rats, p < 0.0001 vs. control group) (Figure 3C and 3D). These findings are comparable to the histological features observed in the MGs from OR patients.

**Figure 3.**
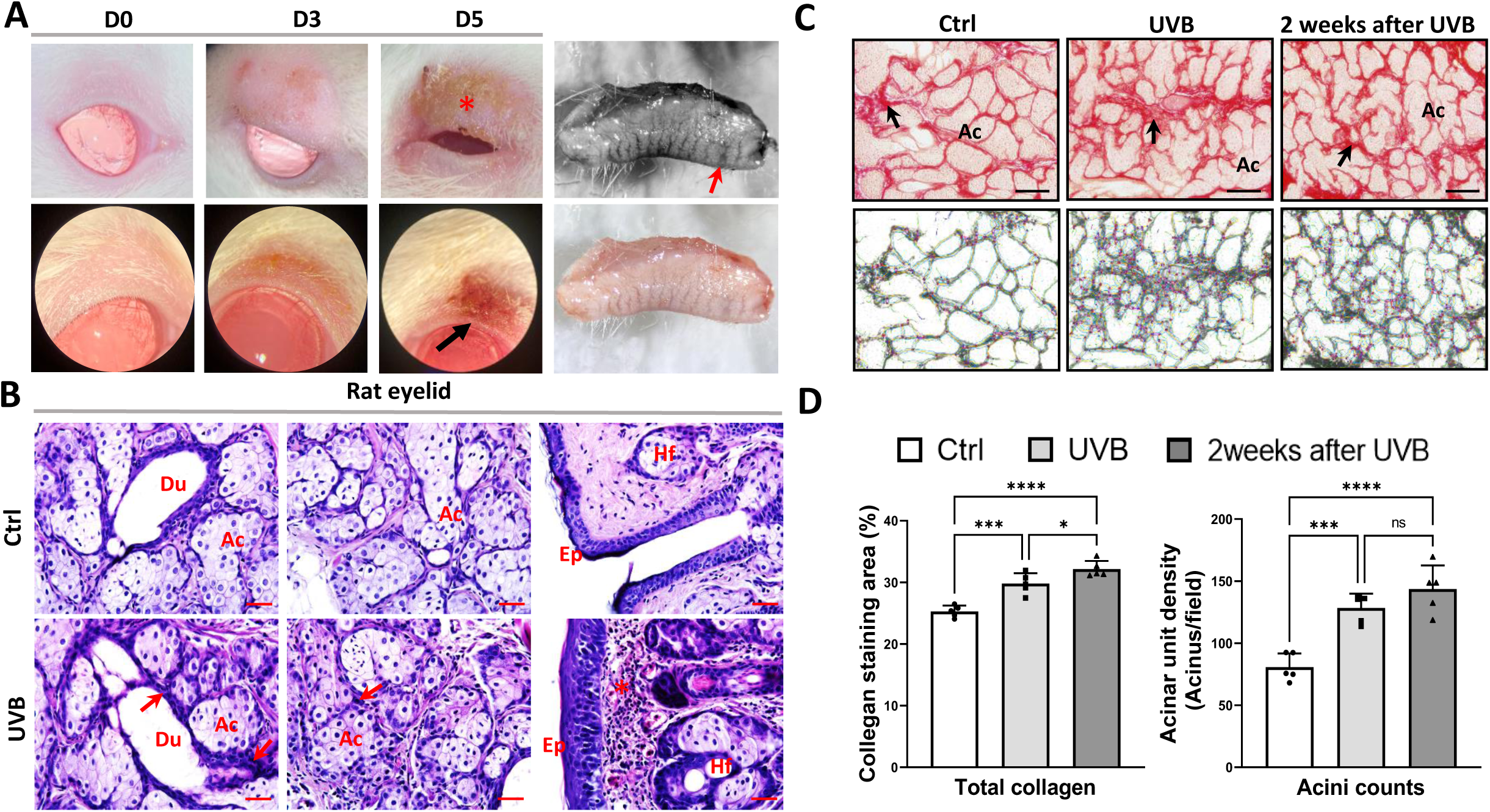
UVB irradiation on rat eyelids causes skin burning and histological changes in rat meibomian glands. **A**. Rat eyelid skin is edematous on day 3 (D3) and covered with yellowish secretions (asterisk) on day 5 (D5) after UVB irradiation. Microscopy shows hyperemia of the eyelid margin (black arrow). The black and white image of the inner side of the rat eyelid shows swelling of the meibomian glands (MGs) (red arrow). **B.** HE staining of rat eyelids shows inflammatory cell infiltration (red arrows and asterisk) around ducts (Du) and acini (Ac) of MGs and beneath the epidermis (Ep) after UVB irradiation compared to the control group (Ctrl). Hf, hair follicles. Scale bar: 50um. **C.** Sirius red stains the collagen in the connective tissue around acini of MGs in the control eyelids (black arrow). The staining is enhanced in MGs after 5 days of UVB irradiation and 2 weeks after UVB exposure (day 19) (black arrows). The acini also appear smaller after UVB irradiation. The lower panels represent the measurement of the density of acinar units using Fiji ImageJ software angiogenesis analysis. Scale bar: 50um. **D.** Quantification of the collagen staining area shows a significant increase in MGs in the UVB group compared to the control group. A further significant increase was observed 2 weeks after UVB exposure. Similarly, the density of acinar units is significantly augmented after 5 days of UVB and 2 weeks after UVB exposure compared to the control group. Data are expressed as mean ± SD, n = 5 rats in each group. **** p < 0.0001, *** p < 0.001, * p <0.05, ns, not significant.

### UVB induced functional changes in rat meibocytes

Nile red detects intracellular lipid droplets and can be used to estimate lipid production. Whilst a strong increase in lipid droplets was observed in the peripheral acinar cells of MGs and in the epidermis after 5 days of UVB irradiation, the staining reduced to the level of the control group 2 weeks after UVB exposure (Figure 4A). Indeed, the surface of Nile red-positive lipid droplets significantly increased on day 5 (6.70 ± 1.12% in the UVB group vs. 2.74 ± 0.31% in the control group, n = 4-7 rats, p < 0.0001), then decreased on day 19, two weeks after UVB exposure (3.83 ± 0.62%, n = 5 rats, p < 0.0001 vs. the UVB group, p = 0.0379 vs. the control group) (Figure 4B).

**Figure 4.**
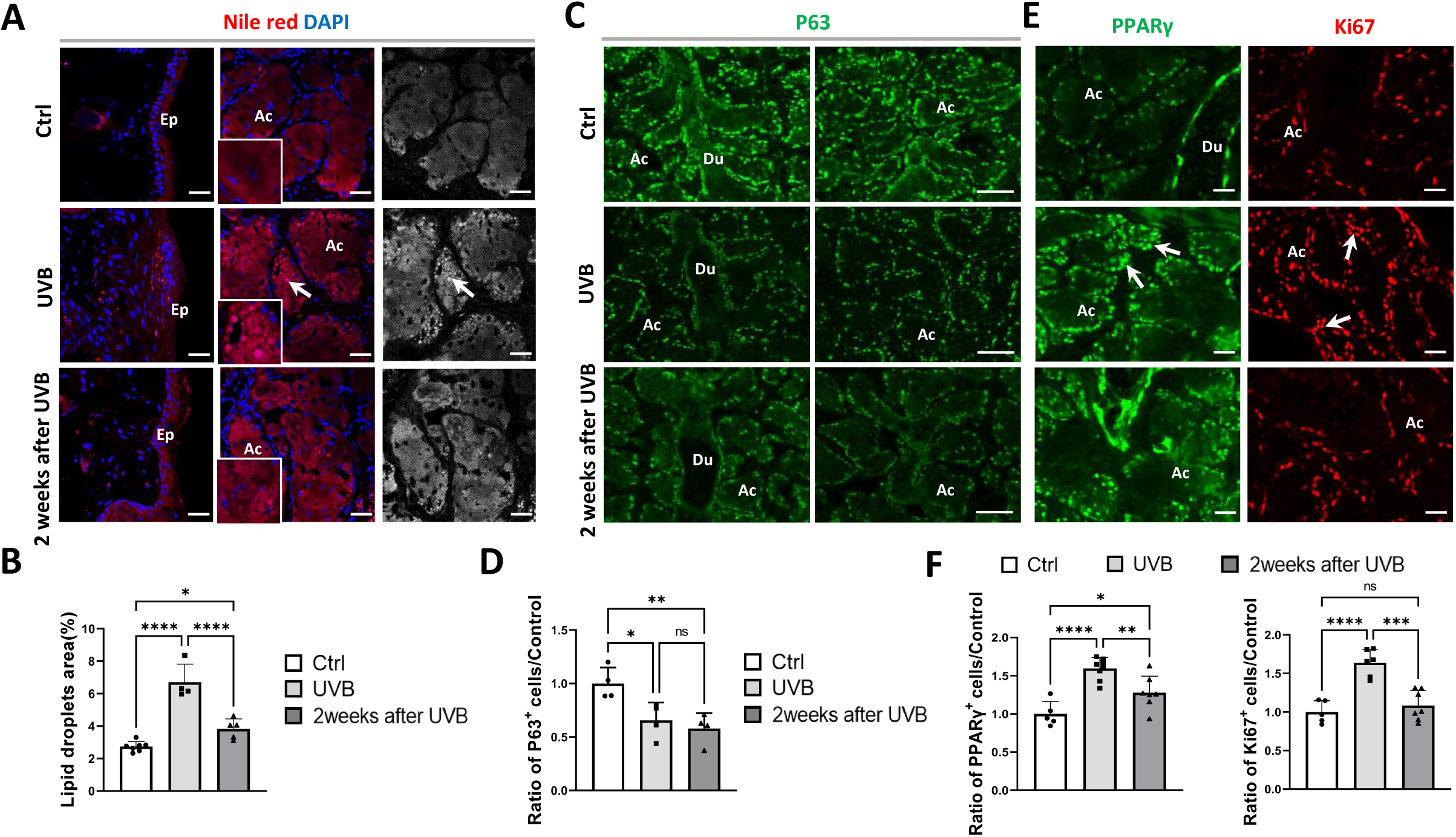
UVB induces functional changes in rat meibomian glands. **A.** Nile red stains lipid droplets (in red) in peripheral acinar (Ac) cells of meibomian glands (MGs) in control rats (Ctrl) without UVB. A diffused staining was also observed in the epidermis (Ep). UVB irradiation enhances Nile red staining in the epidermis and increases lipid droplets (arrows) in acini of the MGs. Two weeks after UVB exposure (day 19), the staining is reduced to a level comparable to the control group. The nuclei are counter-stained with DAPI in blue. Scale bar: 50um. **B.** The area of Nile red-positive lipid droplets is significantly increased in the UVB group compared to the control group. Two weeks after UVB, the lipid droplet area significantly decreased. Data are expressed as mean ± SD, n = 4-7 rats in each group, **** p < 0.0001, * p < 0.05. **C.** P63 immunofluorescence (in green) distributes in the nuclei of epithelial cells of the ducts (Du) and peripheral acinar cells in the MGs of control rats. UVB induces a reduction in P63 staining in MGs. Two weeks after UVB exposure, the intensity and number of P63-positive cells decreases again. Scale bar: 50um. **D.** The ratio of P63-positive cells is significantly decreased in the UVB group compared to the control group. This ratio is even lower two weeks after UVB exposure. Data are expressed as mean ± SD, n = 5-8 rats in each group, ** p < 0.01, * p < 0.05, ns, not significant. **E.** PPARγ (in green) and Ki67 (in red) localize in the nuclei of basal and differentiated acinar cells. UVB irradiation remarkably enhances their expression (arrows) followed by a reduction two weeks after UVB exposure. Scale bar: 50um. **F.** The ratio of PPARγ- and Ki67- positive cells is significantly increased in the UVB group compared to the control group. Two weeks after UVB exposure, their expression ratio reduces to a level comparable to that of the control group. Data are expressed as mean ± SD, n = 5-8 rats, **** p < 0.0001, *** p < 0.001, ** p < 0.01, * p < 0.05, ns, not significant.

P63 is a specific marker of proliferation in stem cells of stratified epithelia [31], and plays essential role in maintaining the integrity and differentiation of acinar cells and epithelial cells lining the ducts of the MG [32]. We found P63 localized in acinar basal cells and in epithelial cells in MG ducts. After UVB irradiation, its expression was progressively reduced from day 5 to day 19 (2 weeks after UVB) (Figure 4C), indicating impaired meibocyte stem cell renewal. The ratio of P63-positive cells to DAPI-stained cells (data normalized to the control group) was significantly decreased after 5 days of UVB exposure (0.66 ± 0.17, n = 5-8 rats, p = 0.0278) and two weeks after UVB (0.58 ± 0.14, n = 7 rats, p = 0.0094 vs. the control group) (Figure 4D).

Ki67 is a marker of cell proliferation, and peroxisome-proliferator-activated receptor γ (PPARγ) plays important role in meibocyte differentiation and lipid synthesis [33]. They are both involved in MG acinar cell renewal and function. PPARγ and Ki67 were expressed in acinar basal cells, peripheral meibocytes, and in epithelial cells in MG ducts. UVB induced a transient increase in their expression on day 5, which is consistent with lipid hyperproduction, followed by a decrease 2 weeks after UVB exposure (Figure 4E). The ratio of PPARγ-positive cells to DAPI-stained cells (normalized to the control group) increased to 1.59 ± 0.14 on day 5 (n = 5-8 rats, p < 0.0001), then decreased to 1.277 ± 0.22 two weeks after UVB (n = 5-8 rats, p = 0.0083 vs. the UVB group, p = 0.0413 vs. the control group). Similar trend was observed for Ki67-positive cells (Figure 4F). The ratio of Ki67-positive cells increased to 1.639 ± 0.17 compared to the control group on 5 days (n = 5-8 rats, p<0.0001), then decreased to 1.083 ± 0.19 (n = 7 rats, p = 0.0001 vs. the UVB group), almost the level of the control group.

### UVB induced hyperkeratinization in rat ductal epithelium and eyelid epidermis

KRT1 and KRT10 are markers for keratinocyte differentiation and cornification [34,35]. KRT10 staining was remarkably enhanced in the ductal epithelium of rat MG after 5 days of UVB compared with the control MG, and its high expression was maintained up to 2 weeks after UVB (Figure 5A). KRT1 expression was particularly increased in the epidermis of rat eyelid after UVB exposure. (Figure 5A). These findings confirmed that UVB induces hyperkeratinization in MG.

**Figure 5.**
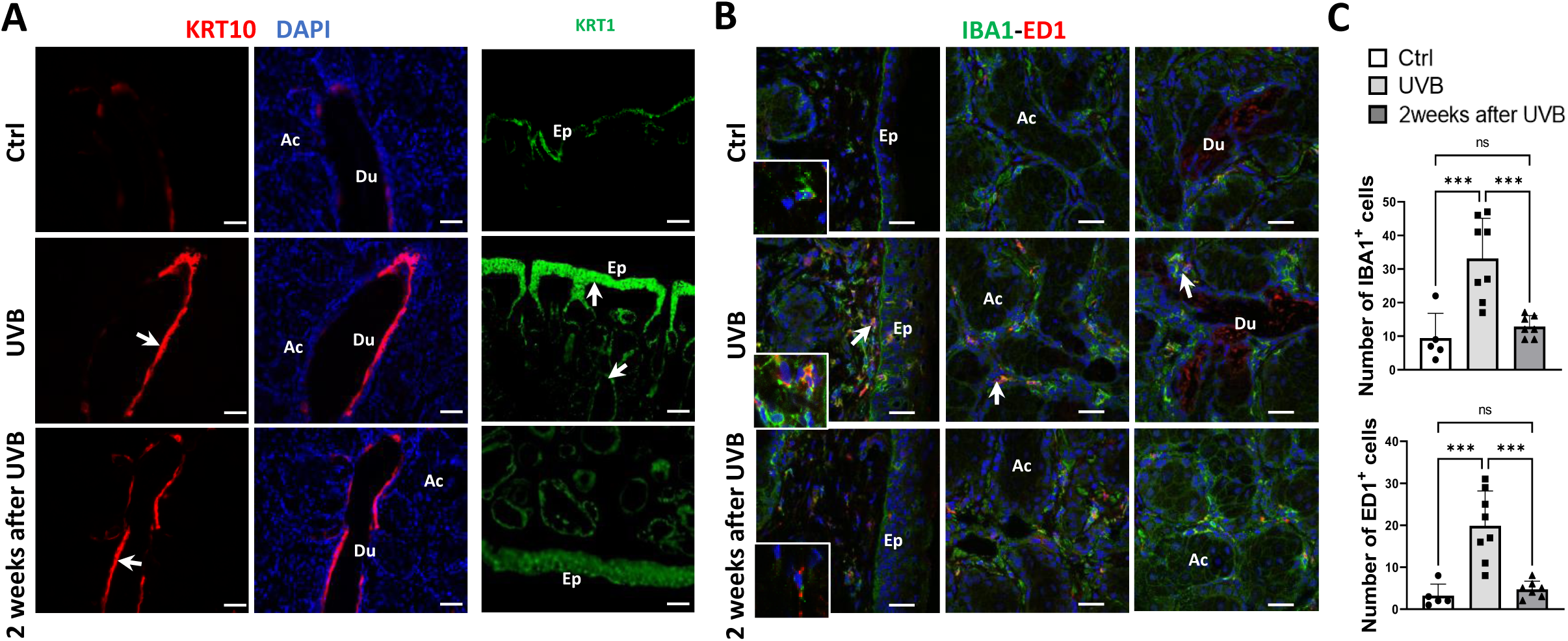
UVB induces hyperkeratinization and inflammatory cell infiltration in rat meibomian glands. **A.** KRT10 (in red) marks slightly the epithelium of the ducts (Du) of meibomian glands (MGs) and KRT1 (in green) labels the superficial epithelial cells of the eyelid epidermis (Ep) in control rats (Ctrl). UVB irradiation enhances the KRT10 expression in ducts (arrow) and KRT1 expression (arrows) in the epidermis. Two weeks after UVB exposure (day 19), the immunofluorescence of KRT10 and KRT1 remains intense. The nuclei are counter-stained with DAPI in blue. Ac, acini. Scale bar: 50um. **B.** IBA1 (in green) and ED1 (in red) label resident immune cells in rat eyelids and MGs in the control group. UVB induces a remarkable increase in inflammatory cells (arrows) infiltrated in connective tissues beneath the epidermis and around acini (Ac) and ducts in MGs. Two weeks after UVB irradiation, IBA1- and ED1-positive cells are reduced. The nuclei are counter-stained with DAPI in blue. Scale bar: 50um. **C.** The number of IBA1- and ED1- positive cells are significantly increased in MGs in the UVB group compared with the control group. Two weeks after UVB exposure, the number of inflammatory cells significantly decreased. Data are expressed as mean ± SD, n = 5-8 rats in each group, *** p < 0.001, ns, not significant.

### UVB induced inflammatory cell infiltration in rat MG

IBA1 and ED1 both stains macrophages in the connective tissues beneath the epidermis, among MG acini and around ducts in the control rat eyelid. UVB irradiation induced a dramatic increase in IBA1- (33 ± 12 vs. 9 ± 7, n = 5-8 rats, p = 0.0005) and ED1-positive macrophages/infiltrated monocytes (20 ± 8 vs. 3 ± 3, n = 5-8 rats, p = 0.0002) in these tissues, suggesting enhanced inflammatory responses. Two weeks after UVB exposure, the number of IBA1- (13 ± 3, n = 7 rats, p = 0.0009) and ED1-labelled inflammatory cells (5 ± 2, n = 7 rats, p = 0.0002) significantly decreased, indicating resolution of inflammation (Figure 5B and 5C).

### UVB induced cell apoptosis, mitochondrial dysfunction, and oxidative stress in rat MG

TUNEL assay revealed the presence of numerous apoptotic cells in UVB-irradiated MGs while the control group without UVB showed barely positive staining (Figure 6A). The TUNEL-positive cells were predominantly located inside and around ducts of the MG, which was consistent with the cell shedding in the ducts observed in the histological sections described above.

**Figure 6.**
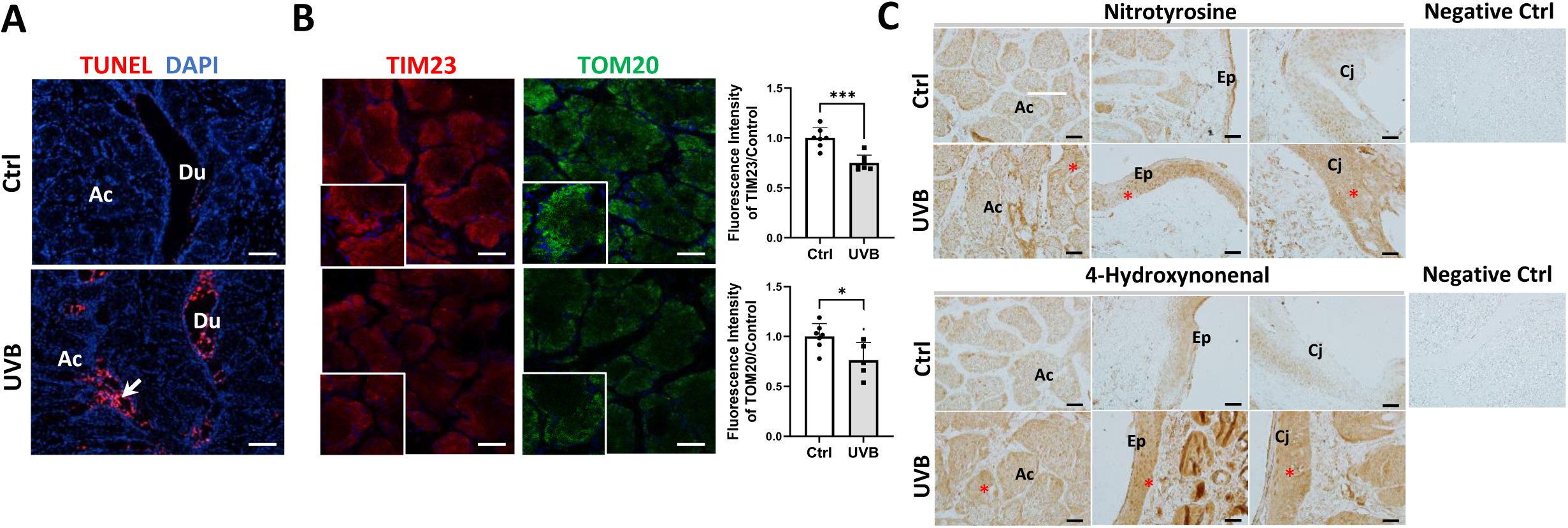
UVB induces cell apoptosis and oxidative stress in rat meibomian glands. **A.** TUNEL assay shows abundant apoptotic cells (arrow) predominantly in the epithelium of the ducts (Du) and the ductules connecting acini (Ac) in the UVB group compared to the control group (Ctrl). The nuclei are counter-stained with DAPI in blue. Scale bar: 50um. **B.** In control rat meibomian glands (MGs), TIM23 and TOM20 distribute in the peripheral acinar cells where mitochondrial activity is high. UVB irradiation considerably reduces their expression. The nuclei are counter-stained with DAPI in blue. Scale bar: 50um. The relative intensity of TIM23 and TOM20 decreases significantly after UVB irradiation. Data are expressed as mean ± SD, n = 5-7 rats, *** p < 0.001, * p < 0.05. **C.** Immunohistochemistry of nitrotyrosine and 4-hydroxynonenal shows slight staining in the acini, epidermis (Ep) and conjunctiva (Cj) of control rat eyelids. UVB irradiation enhances the staining of both markers in all these tissues (asterisks). No staining was detected in negative control. Scale bar: 50um.

TOM20 and TIM23 are markers of the outer and inner mitochondrial membrane complexes essential for mitochondrial function [36]. In the control group without UVB irradiation, TIM23 and TOM20 were distributed in the peripheral acinar cells of the MG. The expressions of TOM20 and TIM23 decreased significantly after UVB irradiation (TIM23: 0.75 ± 0.08, n = 7 rats, p = 0.0002; TOM20: 0.76 ± 0.18, n = 5-7 rats, p = 0.0221), showing that UVB severely affects mitochondrial function (Figure 6B). UVB-induced oxidative stress was evaluated by 3-nitrotyrosine (3-NT), protein nitration marker, and 4-Hydroxynonenal (4-HNE), marker for lipid peroxidation. We observed an enhancement of their expression in the acini of the MG, and in the epidermis and conjunctiva after 5 days UVB exposure (Figure 6C), indicating damage in rat eyelid and MG.

### UVB-induced MGD causes superficial epithelial keratitis

UVB irradiation of rat eyelids induced increased oxidative stress, mitochondrial dysfunction, and apoptosis in MG tissues on day 5 (Figure 7A). Inflammation, hyperkeratinization, meibocyte activity and lipid production peaked on day 5 of UVB exposure, then gradually decreased over the following two weeks (Figure 7B). However, fibrosis in the interstitial tissue continued to increase and proliferation of meibocyte stem cells progressively decreased (Figure 7C), suggesting chronic damage in rat MGs.

**Figure 7.**
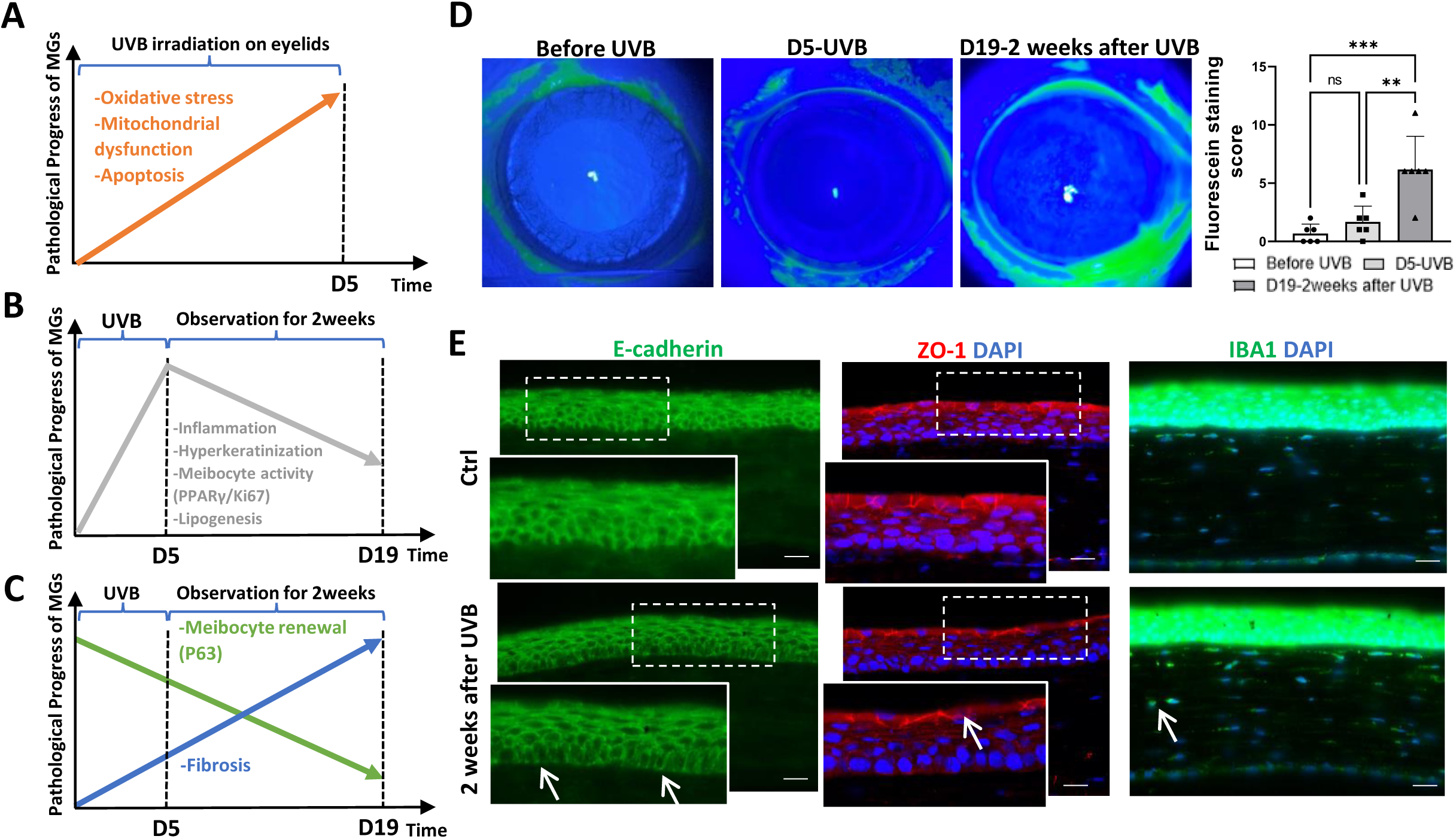
UVB-induced meibomian gland dysfunction leads to corneal defects in rats two weeks after UVB exposure. **A-C.** Kinetics of pathological events observed in rat MGs induced by UVB irradiation. **D.** Slit lamp shows intact corneas before and on day 5 of UVB exposure. On day 19, i.e. two weeks after UVB, fluorescein staining shows diffuse punctuate corneal epithelial defects. Quantification of fluorescein staining score shows significant increase of corneal epithelial defects two weeks after UVB. Data are expressed as mean ± SD, n = 6 rats, *** p < 0.001, ** p < 0.01, ns, not significant. **E.** E-cadherin (in green) homogenously distributes between the epithelial cells of the cornea in control (Ctrl) rats (inset), while its expression reduces two weeks after 5 days of UVB exposure, particularly in the basal cell layer where focal discontinuity can be observed (arrows in inset). ZO-1 (in red) localizes between the superficial epithelial cells of the cornea in the control group (inset). Two weeks after UVB exposure, the superficial layer of the corneal epithelium becomes thinner with focal ZO-1 disruption (arrow in inset). IBA1 stains resident immune cells (in green) in the anterior stroma beneath the corneal epithelium in control rats, whereas active round macrophages (arrow) can be found in the posterior stroma of corneas two weeks after UVB irradiation. The nuclei are counter-stained with DAPI in blue. Scale bar: 20um.

To further investigate the consequences of UVB-induced MGD on the cornea, fluorescein staining was performed before UVB, after 5 days of UVB and 2 weeks after UVB exposure (day 19). As we protected the cornea during UVB exposure, rare fluorescein staining was observed on the cornea after 5 days of UVB. However, two weeks after UVB, diffuse punctate staining was observed on the cornea (Figure 7D). The fluorescein score was 0.67 ± 0.82 before UVB, 1.67 ± 1.37 at day 5 of UVB, and 6.17 ± 2.86 two weeks after eyelid UVB irradiation (n = 6 rats). While no significant difference in corneal fluorescein staining was observed between day 5 and pre-UVB groups, the score increased significantly 2 weeks after UVB exposure (p = 0.0004 vs. the control group, p = 0.0024 vs. the D5-UVB group) (Figure 7D), indicating that the UVB-induced MGD would cause long-term corneal damage.

E-cadherin, a marker of epithelial integrity and cell-cell adhesion was distributed continuously and uniformly between epithelial cells throughout the entire control corneal epithelium. Its expression was reduced 2 weeks after UVB exposure, particularly in the basal layer (Figure 7E, left panels). ZO-1 localized in the superficial layer of the corneal epithelium in the control cornea, was reduced and focally disrupted in the cornea 2 weeks after UVB irradiation (Figure 7E, middle panels). IBA1-positive cells invaded the anterior and posterior stroma (Figure 7E, right panels), suggesting inflammatory responses in the cornea.

### UVB induced changes in the transcriptomic signature of rat MG, comparable to those observed in MGD and rosacea in humans

A bulk transcriptomic analysis was performed to identify the underlying mechanisms involved in UVB-induced MG pathology in rats. Differential express analysis revealed 611 significantly upregulated genes and 660 significantly downregulated genes after UVB exposure (Figure 8A). The heatmap displayed distinct gene expression profiles between the UVB-irradiated and non-irradiated groups (Figure 8B). Over-representation analysis using REACTOME revealed UVB-induced functional gene sets related to keratinization, cornification, mitosis and cell cycle, extracellular matrix organization and degradation, interleukin signaling and complement activation, as well as metabolism diseases (Figure 8C). HALLMARK analysis identified gene sets related to cell cycle checkpoints, epithelial mesenchymal transition, complement, IL6-JAK-STAT3 signaling, inflammatory response, angiogenesis, apoptosis and hypoxia (Figure 8C). GO enrichment analysis highlighted gene sets related to biological processes, including epidermal cell and keratinocyte differentiation, keratinization and cornification, response to oxidative stress, T cell activation, adaptive immune response, lymphocyte-mediated immunity, inflammatory response, leukocyte and granulocyte chemotaxis, DNA damage response signaling, fatty acid biosynthetic and metabolic processes and response to corticosteroid (Figure 8D). Interaction network analysis using GOBP terms (log2FC>1.5) highlighted the most differentially expressed genes including *Ccl2, Defb14, IL-6, Ptgs2 and Ccl7* that are involved in immune regulation, inflammatory responses and chemotaxis, and *Fscn1* that is crucial for cell motility and adhesion (Figure 8E).

**Figure 8.**
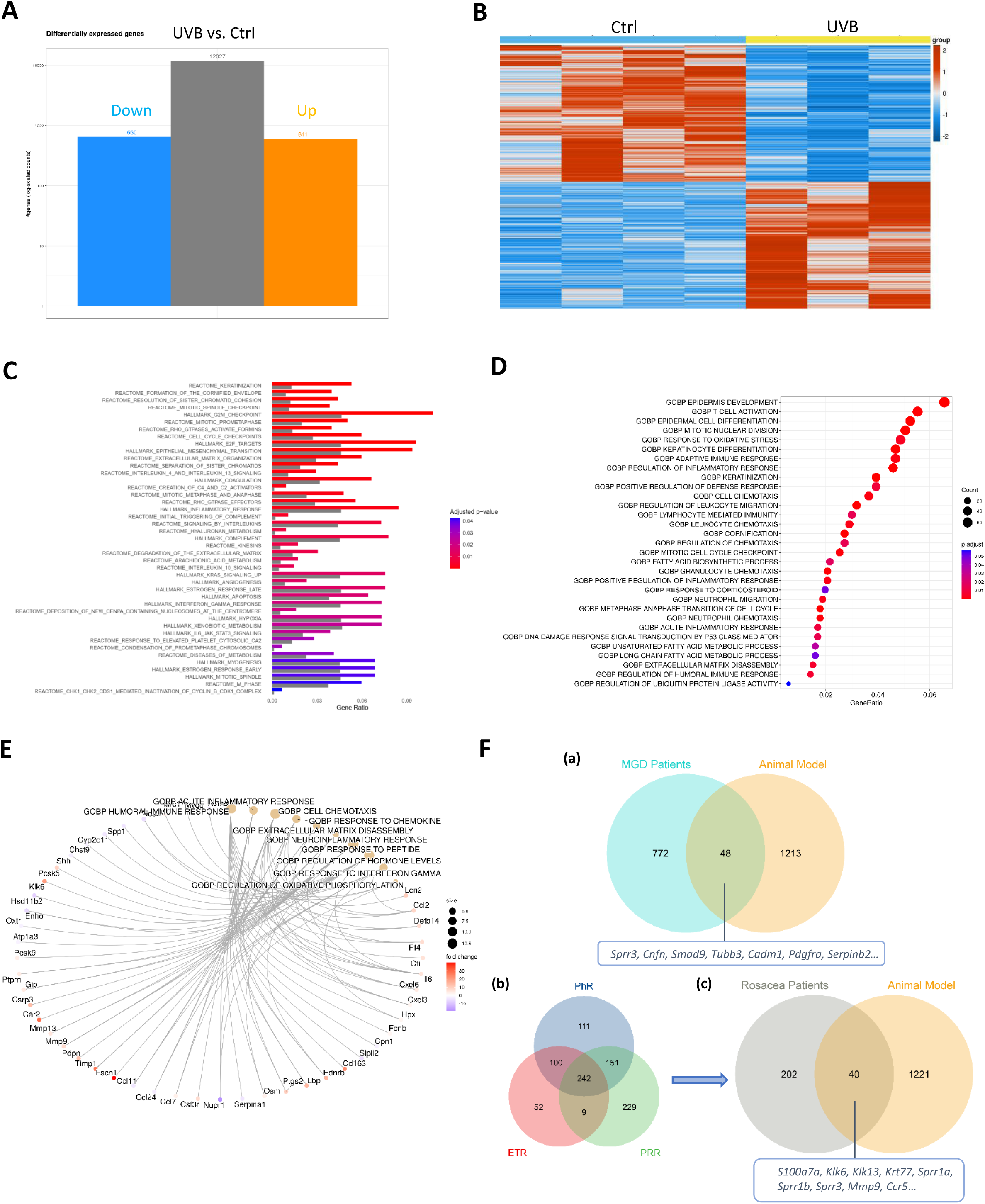
Transcriptomic analysis of UVB-induced changes in rat meibomian glands compared to transcriptomes of human meibomian gland dysfunction and rosacea. **A.** Differential expressed genes plot displayed 611 upregulated and 680 downregulated genes in UVB-irradiated rat meibomian glands (MGs). **B.** The heat map shows distinct gene expression profiles between the UVB-irradiated (n = 3) and non-irradiated groups (n = 4). **C.** Significantly regulated functional gene sets identified by over representation analysis using REACTOME and HALLMARK pathway databases. **D.** Significantly regulated functional gene sets related to biological processes (BP) identified by GO enrichment analysis. **E.** GO network analysis of the top enriched GOBP terms. **F.** Transcriptomic intersection analysis between UVB-irradiated rat MGs and human MG dysfunction (MGD) (GSE17822) highlights 48 commonly regulated genes (a). Transcriptomic analysis of 3 subtypes of the rosacea identifies 242 shared genes (GSE65914) (b). Intersection analysis between rosacea patients and UVB-irradiated rat MGs revealed 40 commonly regulated genes.

Further comparison of MG transcriptomes from MGD patients and UVB-exposed rats identified 48 commonly regulated genes, including *Sprr3*, *Cnfn*, *Smad9*, *Tubb3*, *Cadm1*, *Pdgfra*, *Serpinb2*, and others that play significant roles in the pathogenesis of MGD (Figure 8F, a). The intersection analysis between rosacea patients and the UVB model revealed 40 commonly regulated genes, including especially *S100a7a*, *Klk6*, *Klk13*, *Krt77*, *Sprr1a*, *Sprr1b*, *Sprr3*, *MMP9* and *Ccr5*, which are critical for the development of rosacea (Figure 8F, b and c).

### UVB induced changes in lipid composition in rat MG

The MGs from UVB (n=11) and non-UVB rats (n=17) were subjected to lipidomic analysis. A total of 1406 lipid species from 13 classes were identified: 189 phosphatidylcholines (PC), 136 phosphatidylethanolamines (PE), 104 phosphatidylserines (PS), 67 phosphatidylinositols (PI), 88 phosphatidylglycerols (PG), 51 sphingomyelines (SM), 66 free fatty acids (FA), 66 cholesteryl esters (CE), 113 wax esters (WE), 107 triglycerides (TG), 305 diesters (DE, DE-I and DE-II), 42 ceramides (Cer), and 72 (O-acyl)-ω-hydroxy fatty acids (OAHFA) (Figure 9A).

**Figure 9.**
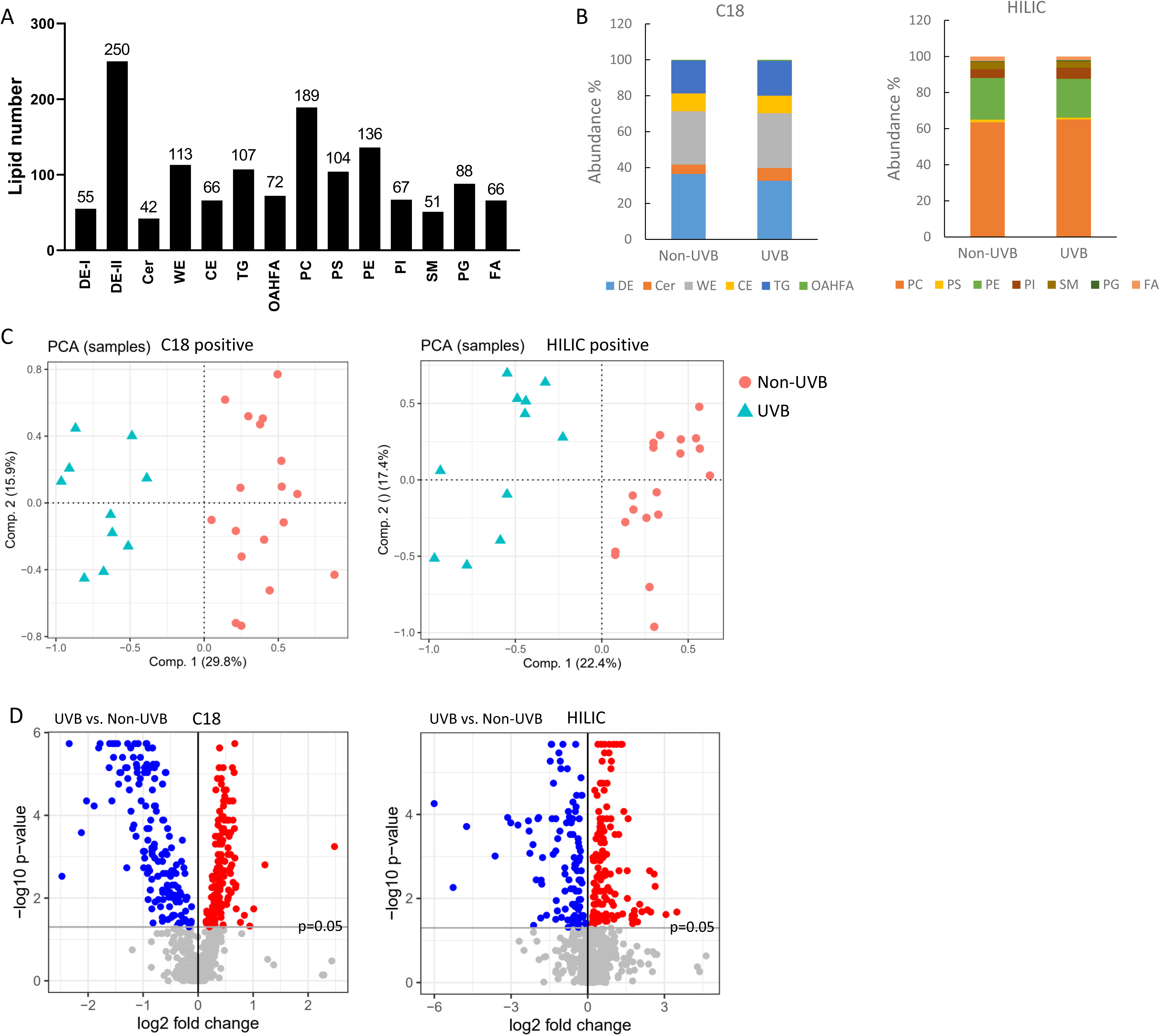
Lipidomic analysis of UVB-irradiated meibomian glands. **A.** Number of lipid species among lipid classes identified in the study. **B.** Proportion of lipid classes in UVB and non-UVB groups. Abundance (%) of lipid classes identified by C18 and HILIC. **C.** PCA analysis illustrates a separation of lipidome signature between UVB and non-UVB groups. **D.** Volcano plots identify lipid species significantly more abundant (in red) and less abundant (in blue) in the UVB group compared to non-UVB group. C18 shows non-polar and polar lipid species with p values < 0.05. HILIC shows polar lipid species with p values < 0.05.

We did not find a significant difference in the proportion of six lipid classes identified by C18 in rat MGs between UVB and non-UVB groups (Figure 9B). However, UVB tended to decrease DE lipids (32.63 ± 7.14% in the UVB group vs. 36.37 ± 2.79% in the non-UVB group, p = 0.063). WE, TG and CE counted for 29-30%, 18-19%, and 9-10% respectively. Cer and OAHFA represented 5-7% and 0.4-0.6% of total lipids identified by C18.

HILIC was used to better detect phospholipid species. Among seven polar lipid classes identified by HILIC, PC were more abundant in the UVB group (65.09 ± 1.70% vs. 63.49 ± 1.44% in the non-UVB group, p = 0.008), while PE (21.50 ± 0.54% in the UVB group vs. 23.00 ± 1.72% in the non-UVB group, p = 0.031) and PS (1.00 ± 0.48% in the UVB group vs. 1.55 ± 0.30% in the non-UVB group, p = 0.008) were less abundant in the UVB group (Figure 8B). The proportion of PI (5-6%), SM (3.5-4%), PG (0.5-0.6%), and FA (2.1-2.6%) was comparable between two groups (Figure 9B).

The positive mode of C18 and HILIC showed clear separation of lipidome signature for the UVB and non-UVB groups on the first axis of the PCA (Figure 9C), while no obvious separation was observed on the PCA map with the negative mode (not shown). This indicates that the main source of variability of the data in positive mode is due to UVB effect. Based on univariate analysis considering both positive and negative modes, the volcano plots showed lipid species that were differentially expressed between two groups (Figure 9D). C18 highlighted 344 lipid species significantly regulated by UVB, among which 178 were more abundant in the UVB group, and 166 were more abundant in the non-UVB group (Supplementary Table 1 and 2). HILIC identified 232 differentially expressed lipid species, among which 97 were more abundant in the UVB-irradiated group and 135 were more abundant in the non-irradiated group (Supplementary Table 3 and 4).

Among the essential non-polar meibum lipid classes, more than half of DE-I (C38-C53), DE-II (C44-C80) and WE species (C34-C54) were differentially expressed (Table 1). Of note, the majority of down-regulated lipid species had greater carbon number while the majority of up-regulated lipid species had smaller carbon number (Supplementary Table 1).

**Table 1.** Number of UVB-induced lipid species in each lipid classes.

|  | Lipid class | N° of total species | N° of up-regulated species | N° of down-regulated species |
| --- | --- | --- | --- | --- |
| Non-polar | DE-I | 55 | 5 | 11 |
|  | DE-II | 250 | 55 | 91 |
|  | WE | 113 | 43 | 29 |
|  | TG | 107 | 17 | 10 |
|  | CE | 66 | 29 | 9 |
| Polar | Cer | 42 | 17 | 2 |
|  | OAHFA | 72 | 12 | 14 |
|  | PC | 189 | 40 | 45 |
|  | PE | 136 | 20 | 51 |
|  | PS | 104 | 3 | 13 |
|  | PI | 67 | 13 | 0 |
|  | PG | 88 | 17 | 0 |
|  | SM | 51 | 3 | 25 |
|  | FA | 66 | 1 | 1 |

Among polar lipid classes, Cer and SM belong to sphingolipid family, UVB induces changes in 19 Cer species (C35-C46) with 17 increased and only 2 decreased, while 28 SM species (C32-C45) were altered with 25 decreased and only 3 increased. Among phospholipids, the majority of PC (C24-C58) and PE species (C29-C46) were differentially expressed between UVB and non-UVB groups. UVB induced minor changes in other phospholipids (Table 1).

## Discussion

The diagnosis of OR is based on non-specific clinical signs, while morphological changes in the MG can be assessed by infrared meibography and other *in vivo* techniques including confocal microscopy and optical coherence tomography [37]. However, the histopathological changes of the MG in OR patients remain largely unknown. In addition to spongiosis, inflammatory cell infiltration and proliferative changes in the eyelid epidermis that have also been observed in skin rosacea [38], we found MG atrophy and fibrosis of the interstitial tissue surrounding the acini in OR patients. The inflammatory infiltrate consists essentially of mononuclear cells, polynuclear neutrophils and eosinophils localized around meibomian acini. In our rat model, UVB induces clinical features and histopathological changes similar to those observed in OR.

Eyelid UVB irradiation causes acute damage to the lid skin and MGs, accompanied by increased oxidative stress, mitochondrial dysfunction, apoptosis, inflammation and elevated lipid production by day 5. During the two-week healing process following the UVB exposure, although inflammation and meibum lipid hyperproduction tend to normalize, MG duct keratinization persists and the depletion of meibocyte stem cells continues, leading to MGD. The progressive fibrosis observed throughout the whole process in rats is comparable to that observed in the MG biopsy tissues from OR patient, suggesting chronic and irreversible damage to the MGs.

MGD is associated with most OR patients. The Meibum, a lipid secretion produced by MGs, forms the outermost layer of the tear film and prevents the evaporation of the aqueous tear film. Any functional changes in MG activity or lipid production can destabilize the tear film and disrupt ocular surface homeostasis, leading to corneal epithelial damage [39]. MGD cans be classified into two forms: the hyposecretory obstructive form, characterized by MG obstruction due to hyperkeratinization of the ductal epithelium [40], and the acinar atrophic form where abnormal meibocyte renewal is the primary cause of MGD [41]. Our finding suggests that UVB exposure to rat eyelid can induce both MG obstruction and acinar atrophy, which may be responsible for the corneal epithelial defects and barrier disruption observed in the later stage.

Transcriptomic analysis of rat MGs and comparisons with transcriptomic data from patients with MGD or rosacea revealed commonly regulated genes involved in the pathogenesis of both conditions. Among these genes, *Sprr1a*, *Sprr1b*, and *Sprr3* encode Small Proline-Rich Proteins (SPRRs), and *Cnfn* encodes Conifelin. Together, these proteins form the cornified cell envelope in terminally differentiating stratified squamous epithelial [42]. The significant upregulation of *Sprrs* and *Cnfn* suggests enhanced keratinocyte and epidermal cell differentiation, which may contribute to hyperkeratinization of the MG ductal epithelium, a primary cause of obstructive MGD [43]. These genes have also been deregulated in papulopustular rosacea [44]. The interaction between keratinocytes and immune cells, mediated by pathways such as Signal Transducers and Activators of Transcription (STATs), is essential in rosacea pathogenesis [45]. Elevated STAT3 activation is also involved in the pathogenesis of MGD-induced dry eye [46]. STAT pathways can be activated by interferons and cytokines such as IL-6 and IL-10, which promote chronic inflammation, tissue remodeling and immune cells activation [47,48]. The S100 protein family, particularly S100A7, S100A8 and S100A9, plays roles in inflammation, immune responses, and cell differentiation [49], and has been implicated in both MGD and rosacea [43,50,51]. In rat MGs, UVB induces an upregulation of *S100a7* and a tendency in increasing *S100a8* and *S100a9,* which may be involved in inflammatory processes in MG tissues. Significant activation on cell chemotaxis is involved in the adaptive immune responses in rosacea patients [50]. CCL2 and CCL7 attract monocytes, macrophages and dendric cells and bind to receptors such as CCR2, guiding their migration to inflammatory sites [52]. These chemokines recruited cells release cytokines including IL-6, which further amplifies the inflammatory response by promoting T cells and B cells differentiation and enhancing CCL2 expression [53,54]. This is particularly relevant in chronic inflammatory diseases like rosacea, where persistent recruitment of monocytes and macrophages exacerbates tissue damage [50]. SERPINB2, also known as plasminogen activator inhibitor type 2 (PAI-2) is widely expressed in the epithelium (including in the conjunctiva and cornea), keratinocytes, fibroblasts and macrophages [55], and is involved in extracellular matrix (ECM) organization, inflammation and adaptive immune response [56,57]. *Cadm1* is downregulated in both UVB-exposed rat MGs and human MGD [58]. CADM1 is expressed in keratinocytes, vascular endothelial cells, neurons as well as immune cells including dendritic cells, T cells, NK cells and mast cells. CADM1 on keratinocytes acts as a scaffolding molecule for immune cells; CADM1 on vascular endothelial cells promotes the repair of the endothelial barrier; and CADM1 on immune cells contributes to the regulation of innate and adaptive immune responses [59,60]. MMPs, upregulated in both rat MGSs and rosacea patients, are involved in the degradation of the ECM. The decreased cell adhesion and increased ECM degradation facilitate cell migration and epithelial-mesenchymal transition, contributing to tissue remodeling. Genes encoding fibronectin and collagen 1, associated with fibrosis, are also regulated by UVB.

Genes involved in the cell cycle and DNA repair are also commonly upregulated by UVB exposure and human MGD [43]. Cyclin-dependent kinases (CDKs) and cyclins promote cell division and MG regeneration [61]. However, abnormal differentiation and senescent cell accumulation contribute to MGD, by disrupting lipogenesis and increasing inflammation [62]. Upregulation of genes related to DNA repair suggests active cellular processes to restore normal gland function in response to UVB-induced DNA damage in the MG [43].

The lipid composition of rat MGs has not been studied previously. We used a non-targeted approach by coupling liquid chromatography and high-resolution mass spectrometry that allows to identify over 1400 lipid species in 13 classes, more than those found in humans (614) [63] and mice (1089) [64] with similar analysis approaches. In human meibum, non-polar lipids including WE, CE, DE and TG are predominant lipid classes that form the lipid layer of the tear film and prevent tear evaporation [65,66]. Although polar lipids secreted by MG including phospholipids, sphingolipids, and amphiphilic OAHFA are minimal, they spread in the interface of the lipid and aqueous layers of the tear film and play an essential role to stabilize the tear film [66,67]. In our study, UVB exposure does not significantly alter the proportion of non-polar lipid classes, but does affect phospholipids. We identify PC as the most prominent phospholipids in rat MG and UVB increases PC including PC 34:1 and PC 34:2, the most abundant PC species. This result is consistent with a former study of human Meibum, in which PC was the most abundant phospholipid class and PC 34:1 and PC 34:2 were consistently detected. In this study, PC level was elevated in patients with dry eye disease [68].

At individual lipid species level, UVB induces significant changes in meibomian lipid profile. Among non-polar lipids, over half of DE, WE, CE lipid species are differentially expressed with a decrease in long chain and extremely long chain lipids essential for tear evaporation resistance [69]. In a pilot study comparing non-polar lipids in Meibum from patients with and without dry eye, many species of DE-I, DE-II and extremely long chain WE are decreased in patients with dry eye [70]. Some of these lipid species have also been identified in our study, including WE 48:2, WE 45:2, WE 45:3, WE 47:2, DE-I 49:2, DE-II 64:3, DE-II 64:4, DE-II 65:2, DE-II 65:3, DE-II 66:3, DE-II 66:4, DE-II 67:2, DE-II 67:3, DE-II 68:3, DE-II 68:4, DE-II 69:3 and DE-II 70:4 that are decreased in the UVB group. It has also been reported that changes in abundance of some lipid species are specific to clinical features of MGD [71]. Among lipid markers associated with eyelid margin hyperaemia, eyelid margin irregularity and MG orifice plugging, we found that CE18:1, CE20:1, CE20:2, CE22:3, CE16:0 and CE18:2 are increased by UVB, and TG55:3 is decreased by UVB. CE30:1, the most abundant CE in Meibum, that has been shown to decrease with age in a mouse model of age-related MGD [64], is less abundant in the UVB group.

Cer, SM are polar lipids belong to the sphingolipid family. *In vitro* and animal studies have found increased Cer to be associated with tear film instability and clinical findings of MGD [72,73]. Sphingolipids composition has also been found to differ in individuals with various dry eye and MGD features [71,74,75]. Although UVB does not induce significant differences in total Cer and SM composition in rat meibum, the majority of differentially expressed Cer species are up-regulated by UVB, while the majority of differentially expressed SM species are down-regulated by UVB. Of note, Cer and SM are also bioactive lipids that regulate important cellular functions such as apoptosis and inflammation. Under stress conditions, such as UVB irradiation [76], Cer production is activated through SM hydrolysis to ensure a fast response to stimuli. The higher abundance of certain Cer species and lower abundance of certain SM species may reflect the cellular production of Cer in the MG induced by UVB. Furthermore, increased abundance of Cer, especially those with long chain may destabilize the tear film leading to rat cornea defect.

UVB-induced morphological and functional changes in rat MGs, along with alterations in meibomian lipid composition, resemble those seen in human MGD and rosacea.

We recognized several limitations. The UVB energy used to irradiate rat eyelids in this study was around 3-5 times higher than the minimal erythema dose for skin [77]. But we chose this dose to ensure sufficient inflammatory response in the MG located in the posterior eyelid. Although it did not cause any eyelid skin ulceration, the UVB dose and the regimen of exposure could be optimized in the future. The long-term effects of UVB exposure on MG function and Meibum lipid composition have not been addressed in this study, which could be further investigated.

## Conclusion

UVB irradiation of rat eyelids induces morphological and functional changes in the MG and subsequent ocular surface damage, comparable to those observed in human OR. UVB-induced changes in lipidomic and transcriptomic profiles in rat MG are similar to human MGD and rosacea. This model can be valuable for studying the pathophysiology of MGD associated with OR.

## Supporting information

Transcriptome of UVB-induced MGD and OR model on eyelids_differetial expressed genes

Intersection analysis for transcriptome of human MGD and UVB-induced model

Intersection analysis for transcriptome of human Rosacea and UVB-induced model

## Acknowledgments

The authors thank the Functional Exploration Center (CEF) and the Core Facilities of Histology, Imaging and Flow Cytometry (CHIC) at Centre de Recherche des Cordeliers.

## Author Contributions

Conceptualization, F.B.-C. and M.Z.; data curation, L.Z. and M.Z.; formal analysis, L.Z., M.C., C.P. and M.Z.; funding acquisition, M.Z. and F. B.-C.; investigation, L.Z., E.G., D.R.-B., M.C., C.L., S.M., X.M., J.-L. B. and M.Z.; methodology, L.Z., D.R.-B., M.C., O.B. and M.Z.; project administration, F.B.-C. and M.Z.; resources, J.-L.B., O.B., F.B.- C. and M.Z.; supervision, O.B., F.B.-C. and M.Z.; validation, F.B.-C. and M.Z.; writing-original draft, L.Z., M.C., C.P. and M.Z.; writing-review and editing, F.B.-C., M.Z. and L.Z..

## Data Availability Statement

All Data are available upon reasonable request. Raw RNAseq data will be made available on Gene Expression Omnibus GSE277020 (accessible on 11 March 2025).

